# Compartmental and spatial rule-based modeling with *Virtual Cell* (VCell)

**DOI:** 10.1101/146225

**Authors:** M. L. Blinov, J. C. Schaff, D. Vasilescu, I. I. Moraru, J. E. Bloom, L. M. Loew

## Abstract

In rule-based modeling, molecular interactions are systematically specified in the form of reaction rules that serve as generators of reactions. This provides a way to account for all the potential molecular complexes and interactions among multivalent or multistate molecules. Recently, we introduced rule-based modeling into the Virtual Cell (VCell) modeling framework, permitting graphical specification of rules and merger of networks generated automatically (using the BioNetGen modeling engine) with hand-specified reaction networks. VCell provides a number of ordinary differential equation (ODE) and stochastic numerical solvers for single-compartment simulations of the kinetic systems derived from these networks, and agent-based network-free simulation of the rules. In this work, compartmental and spatial modeling of rule-based models has been implemented within VCell. To enable rule-based deterministic and stochastic spatial simulations and network-free agent-based compartmental simulations, the BioNetGen and NFSim engines were each modified to support compartments. In the new rule-based formalism, every reactant and product pattern and every reaction rule are assigned locations. We also introduce the novel rule-based concept of molecular anchors. This assures that any species that has a molecule anchored to a predefined compartment will remain in this compartment. Importantly, in addition to formulation of compartmental models, this now permits VCell users to seamlessly connect reaction networks derived from rules to explicit geometries to automatically generate a system of reaction-diffusion equations. These may then be simulated using either the VCell partial differential equations (PDE) deterministic solvers or the Smoldyn stochastic simulator.

## Introduction

The specification of all molecular species and interactions is usually the first step in modeling a biomolecular interaction network. However, for the interactions of multivalent or multistate molecules, the number of species and reactions can be combinatorially large (1, 2), making it impractical to specify the reaction network manually. Rule-based modeling (2, 3) overcomes this limitation by accounting for the complete set of reactions and species that arise when an initial (seed) set of species is transformed using reaction rules. The reaction rules can serve either as generators of individual reactions, expanding the initial set of species into the complete network of reactions and species, or as generators of stochastic events, producing molecular complexes with non-zero population numbers.

Virtual Cell (VCell, http://vcell.org) is an open-source platform that provides powerful capabilities for kinetic modeling of cellular systems (4, 5) (Fig. 1). A key focus of VCell is to allow modelers to ask how spatial features of cells affect the system behavior. At the simplest level, the relative sizes of compartments affect the concentrations of species transported between them; models that account for the surface areas of membranes and the volumes of volumetric compartments, but assume that diffusion is fast on the timescale of reaction kinetics, will be referred to as ‘compartmental’. If diffusion and spatial localization of molecular species can affect the biology, the geometric shapes of the membrane and volumetric compartments also need to be explicitly considered, and these models are considered ‘spatial’.

**Fig 1.**
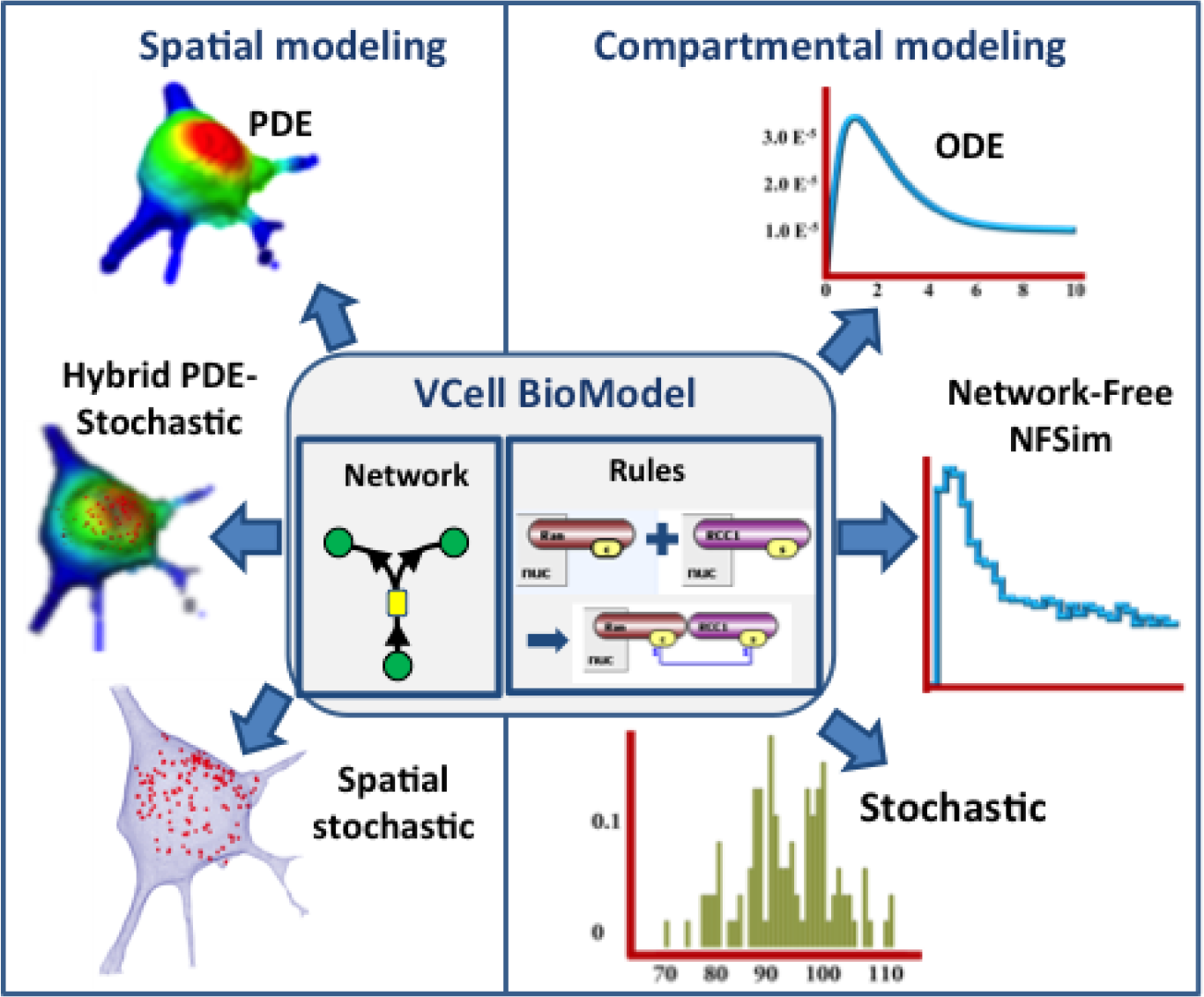
Overview of VCell capabilities and v. 6.1 enhancements. The VCell BioModel (at the center) can be defined as an explicit reaction network, a set of reaction rules, or as a combination of the two. Before VCell 6.1, only single-compartment models defined with reaction rules could be simulated, which also restricted rule-based modeling to non-spatial Applications. VCell 6.1 now supports rule-based modeling with all these VCell solvers. In addition, a new compartmental NFSim solver is introduced.

In building a VCell ‘BioModel’, users initially describe the system ‘Physiology’. with compartments defined as multiple volumes (e.g. extracellular space, cytosol, nucleus, endoplasmic reticulum, etc.) and surfaces (e.g. plasma membrane, mitochondrial membrane, etc.); the ‘Physiology’ also encompasses reactions and fluxes taking place within and between volumes and surfaces, with respective volume-and area-based units for concentrations and kinetic rate expressions. Once a ‘Physiology’ is defined in VCell, any number of ‘Applications’ can be defined, which specify the initial conditions, compartment sizes and/or geometries, and whether the system should be treated deterministically or stochastically. An ‘Application’ can be considered to be a virtual experiment and is sufficient to completely define the system’s mathematical equations, which are automatically generated. VCell applications that do not include explicitly defined geometry (i.e. making the assumption that diffusion is fast on the timescale of all reaction kinetics) are called *compartmental*. These can be simulated using a variety of deterministic and stochastic numerical solvers (Fig. 1) to produce timecourses for concentrations and/or population numbers of species. Alternatively, an explicit geometry can be defined using a variety of methods, such as analytic equations in 1, 2 or 3D, constructed solid geometry in 3D, image-based (imported from various microscopy formats), mesh-based (imported from STL image), or drawn using provided drawing tools. Such VCell applications are called *spatial*, and diffusion for species must be defined in order to simulate timecourses. Diffusion can be defined in both volumetric compartments and along the surface of membranes, and reactions spanning multiple compartments account for species flux between these compartments. VCell offers several solvers for partial differential equations (4–7) to simulate spatial and temporal changes in concentration when the species copy numbers are large. When the copy numbers are low, a spatial stochastic simulator using the Smoldyn simulation engine (8) is available. VCell even has a solver for spatial hybrid deterministic/stochastic simulations (9), to accommodate systems containing some species at high copy number (modeled as continuous concentrations and partial differential equations) and others at low copy number (modeled with stochastic reaction kinetics and Brownian motion).

Earlier versions of VCell required explicit specification of species and reactions. Last year we introduced VCell 6.0, which incorporated a graphical user interface (GUI) to represent multiple sites and states within molecules and the rule-based reaction kinetics between them (10). This GUI provides a compact method for describing the key structural features of multivalent multistate molecules that control their roles in complex signaling systems. Every chemical species can be represented as structured objects composed of molecules, with reactions that control all their modifications and changes in their connectivity. Every model can be simulated both as a reaction network (following network-generation using the BioNetGen engine), permitting both deterministic and stochastic simulations, or with the NFSim network-free algorithm, which produces stochastic simulations. However, the abstractions within the representations of molecules and rules, as well as the algorithms within the network generation and simulation engines did not include compartments or the ability to simulate reaction diffusion equations in explicit geometries.

In this paper we describe a compartmental extension of the rule-based modeling capabilities of VCell, available in VCell 6.1. It enables specification of the locations of molecules and rules. This required us to develop new abstractions to anchor molecules explicitly to volume or surface compartments. We then modified the BioNetGen code to support network generation within the VCell compartmental formalism. This permitted us to support all the stochastic or deterministic, and non-spatial or spatial, simulators available in VCell to simulate reaction networks generated by rules (Fig. 1). Additionally, we modified the NFSim code (11) to support compartmental (albeit non-spatial) network-free simulations.

## Results

Rule-based modeling in VCell is implemented by adapting the standalone software tools BioNetGen (3, 12) and NFSim (11). They allow both deterministic and stochastic simulations after the reaction network is generated (BioNetGen) and network-free particle-based simulations (NFSim) in a single compartment. Both tools operate using the BioNetGen Language, BNGL (12), which was originally designed to work only for a single compartment.

A compartmental extension of BNGL, cBNGL (13), enables explicit modeling of the compartment topology of the cell and its effects on system dynamics using the BioNetGen network generation engine. However, it is not suited for VCell. The major reason is that cellular topology in cBNGL is restricted to a compartment graph in which nodes represent compartments and directed edges represent containership. This graph must be a tree, a membrane may contain (and be contained in) only a single volume compartment, while a volume compartment may contain multiple surface compartments but be contained only by a single membrane. This cellular topology is more limited than the generalized topology available in VCell, where no restrictions are imposed on how compartments can be enclosed within each other. Likewise, cBNGL does not allow for molecular species to span several membranes. The VCell paradigm gives the modeler more flexibility and supports representation of multicellular structures with gap junctions or tight junctions. Additionally, NFSim does not support compartmental simulations using cBNGL. Therefore, to enable rule-based modeling in a generalized topology using both the BioNetGen and NFSim engines, we decided to develop our own schema based on the VCell multi-compartment reaction diagram, where the locations of each reactant, product, and of the reaction itself are explicitly specified. This required new conventions for specifying spatial features within a regular BNGL file. We use the BioNetGen or NFSim engines to generate the reactions, but have added a processing step after every iteration to fix the location of each newly generated species and remove invalid species and reactions. While the BNGL is hidden to the user, it is important to lay out the key algorithmic features that allow for merging of rule-based modeling within the VCell architecture. We describe these in Table 1 and Figure 2, which also serve as a short glossary of rule-based modeling terminology. Note that while a species is located in a given compartment, its orientation in space (as may be required for agent-based simulations) is implicitly determined by the specification of binding reaction rules between sites on molecules with different locations. Supplemental material (S1.pdf) discusses implementation in more detail.

**Table 1.** Concepts and definitions for compartmental rule-based modeling in VCell

| Term | Purpose | Composition | Compartmental/Spatial features |
| --- | --- | --- | --- |
| Compartment | Cellular structure containing species and reactions | A volume or a surface, corresponding to extracellular regions, cytosol, organelles and their associated membranes. No specification of the relationship between compartments (such as enclosures or adjacency) is required. | For **Applications** without explicit geometries, compartmental sizes are specified as surface areas or volumes. For spatial **Applications**, compartments may be explicitly associated with regions within a geometry. |
| Molecule (Fig 2A). | A building block for species. | Comprised of sites that can bind other sites between or within molecules. A site may also have multiple states. In this way, a molecule spawns a collection of chemical species – one per every combination of site occupancy and/or state. | Can be, optionally, **anchored** to one or more compartments. A species containing an anchored molecule can be located only in one of these compartments. |
| Species (Fig 2B). | An individual chemical species that may occur in the model. | Comprised of molecules that are connected through bonds between binding sites. All modification sites must be explicitly defined. A species can be a **seed species** (defined as initial condition and used as a seed for reaction rules application) or a generated species (a result of reaction rule application). | Every species is located in a unique compartment. Seed species are assigned to a compartment by the user. It may not contain molecules that are excluded from that compartment by anchoring to other compartments. |
| Pattern (Fig 2C). | Specifies a set of possible states of species to be selected as participants in reaction rules and in observables. | Comprised of molecules. The states of sites may be left unspecified; thus a pattern may select multiple species. Moreover, binding sites may have **implicit binding status** (*has external bond or may be bound*) where its binding partner is not explicitly defined. Such patterns may be inclusive of species that contain molecules not explicitly specified in a pattern but being possibly bound to molecules within it. | Defined in a single compartment; all molecules that comprise a pattern must be permitted to be located in this compartment. |
| Observable (Fig 2D). | Specify simulation outputs of interest. | Consists of one or more patterns that define features of species. The result is the total population (concentration or count) of multiple species. | Defined in a single compartment; all molecules that comprise an observable must be permitted to be located in this compartment. |
| Reaction rule (Fig 2E). | Defines transformation of multiple species at once, generating multiple reactions | Species to be transformed are selected by reactant pattern(s). Product pattern(s) define the end result of transformation. Product may differ from reactant by re-assigning molecules, adding or deleting bonds and changing site states. A kinetic expression is also a component of a reaction rule. | Reaction rule is defined in a single compartment. Each reactant pattern and each product pattern are assigned a specific compartment, which may be different from compartment for reactants or products. |

**Fig 2.**
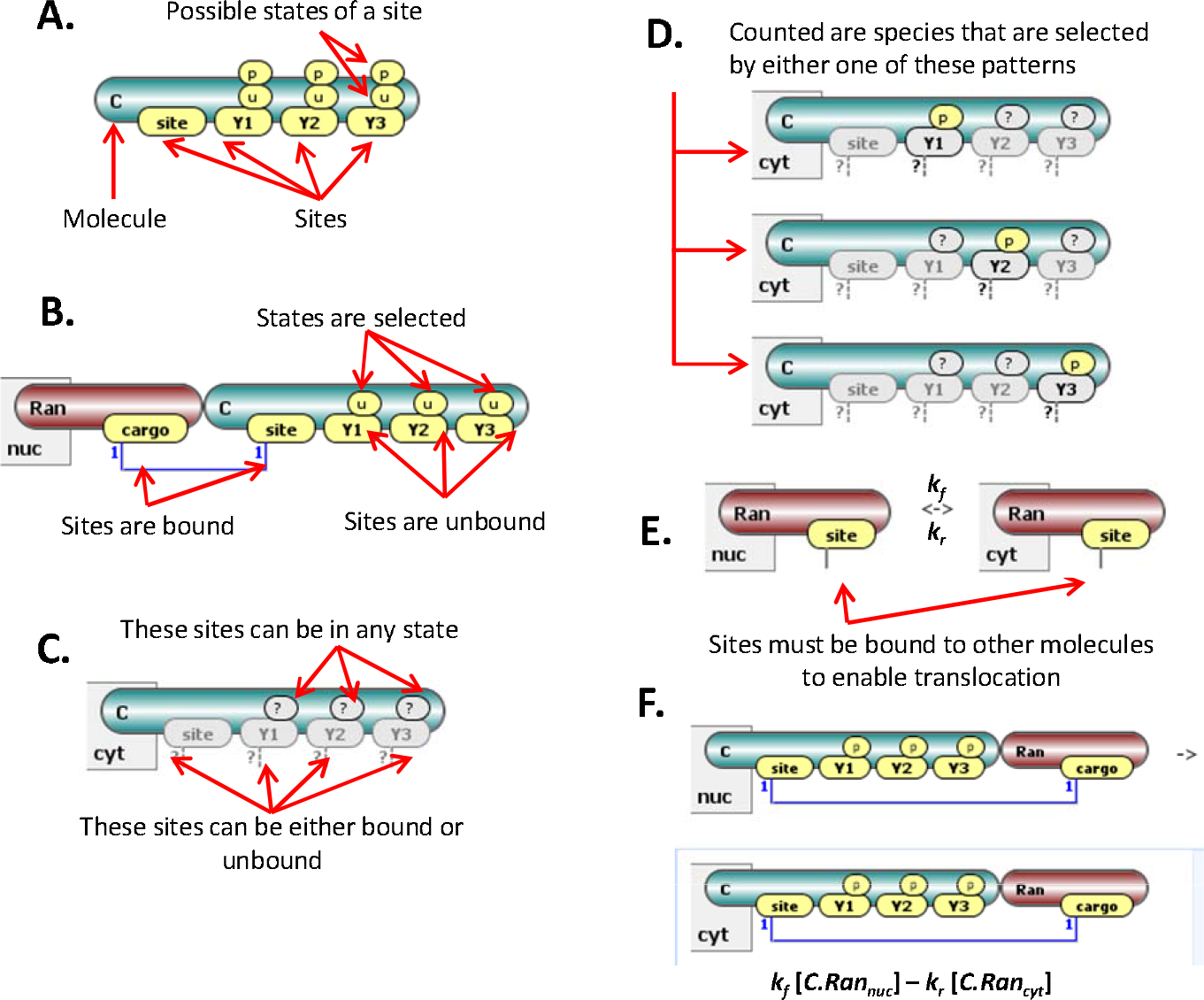
Elements of compartmental rule-based modeling. **A**. Cargo molecule C with three tyrosines subject to phosphorylation and one extra binding site. This molecule spawns at least 8 different potential species – combinations of possible states of phosphosites. **B**. A species – a complex of Ran and C1 in the nuc compartment. All three tyrosines of C are unphosphorylated. **C**. A pattern that includes the set of all cytosolic species that have C as its constituent. **D**. An observable that counts all species that have at least one phosphorylated residue of C among its constituents. **E**. A reaction rule for Ran translocation between nucleus and cytosol. The solid line emanating from the site specifies that Ran must be bound to other molecules to translocate. This is a convenient feature when multiple cargo molecules can be bound to this site – there is no need to enumerate all of them. The rate for all reactions generated by the rule is given by the same rate parameters *k_f_* and *k_r_*. **F**. A sample reaction generated by the rule is parameterized by the same rate constants as the rule.

Let us illustrate compartmental rule-based modeling using a model of Ran-mediated nucleocytoplasmic transport (available in the VCell Database under Tutorials as “Rule-Based_Ran_Transport”). This is a simplified version of a published model by Smith et al. (14). In this tutorial (Fig 3A), the nuclear Ran binds to a cargo molecule, facilitating translocation into the cytosol. Ran and cargo then dissociate. The cytosolic cargo molecule may be phosphorylated on any of its three tyrosines while in the cytosol. When cargo is displaced by the Ran exchange factor RCC1, which is a component of histones, Ran stays in the nucleus. The rule-based approach provides a compact way of describing such a system, with a single transport reaction rule (Fig 2E and #7 in Fig. 3A) in place of 18 reversible reactions for transport of multiple types of cargo. The total number of species and reactions, if all combinations of phosphoforms are accounted for, would be 36 and 98 respectively. Such a large system is difficult to construct, visualize and analyze without rule-based specification (Fig 3B). Moreover, model predictions have to be compared to experimental observations that often correspond to sums of multiple species. In our example, the total phosphorylation of cargo is summed over all species that have at least one site being phosphorylated, with triple-phosphorylated species counted three times. The observable concept (Fig. 1D and Table 1) introduced in rule-based modeling provides an easy way to define such quantities.

**Fig 3.**
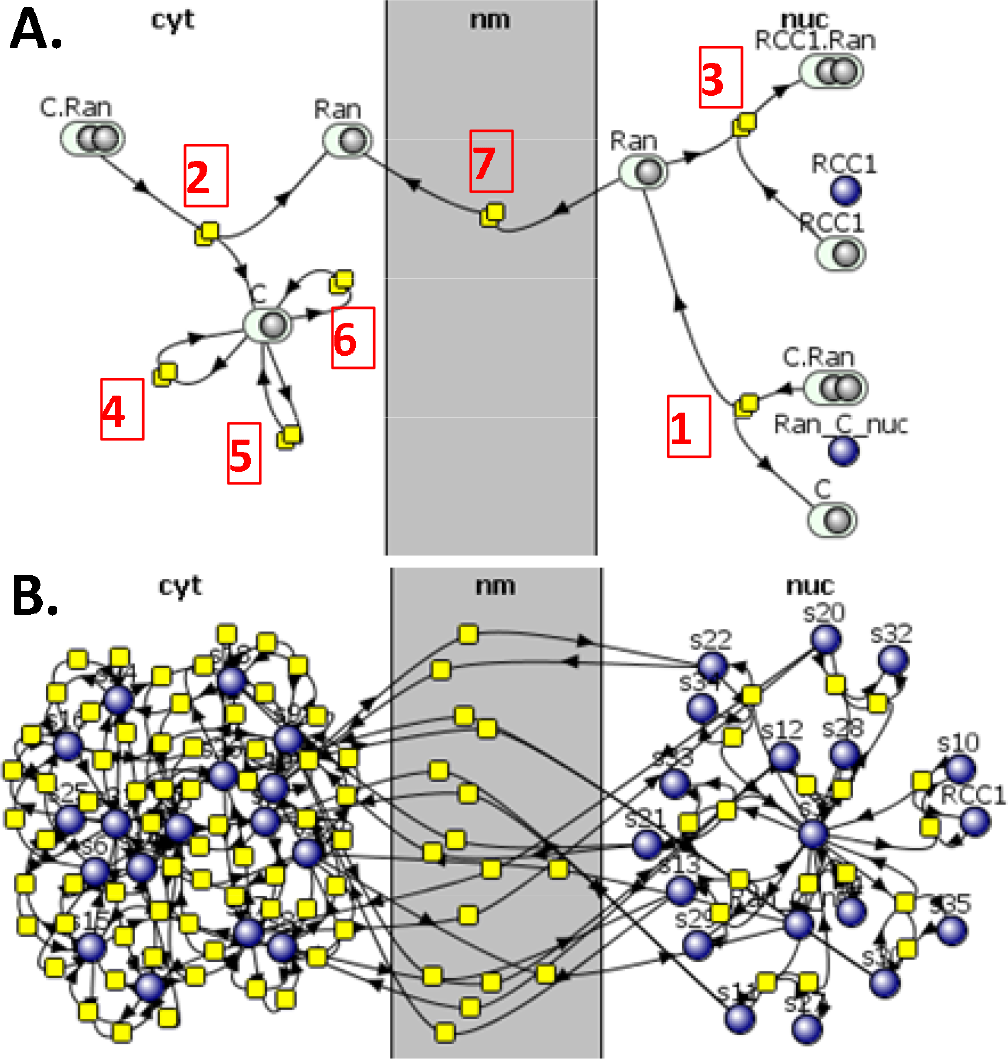
Graphical representation of reaction rules and reaction networks. **A**. A collapsed VCell reaction diagram of a rule-based model for Ran transport across nuclear membrane. Ran can bind cargo molecule C in both nucleus (rule 1) and cytoplasm (2). In nucleus Ran can bind RCC1 (3), while in cytoplasm cargo protein may undergo phosphorylation on all their phosphosites (4–6). Ran can undergo transport across the nuclear membrane (7). The full mechanisms (such as what should be bound to Ran in rule 7) are hidden in this view, but can be seen when the user clicks on a reaction rule node. **B**. The reaction network generated within VCell from this set of rules contains 36 species and 98 reactions.

A feature of rule-based modeling is that anything not explicitly constrained is allowed. Thus, a transport reaction rule for Ran with a pattern specifying it must have a bound site (Fig 2E), will carry with it any of the molecules that may be bound to it, including RCC1, which should remain in the nucleus according to the known biology. To address this within the generalized topology of VCell, we introduced the ability to *anchor* molecules to compartments (Fig 4). A molecule can be anchored to one or multiple compartments. Anchoring a molecule to specific compartments constrains any species (produced through network generation or by NFSim) that contains this molecule to be in only these compartments. In spatial applications, such species are free to diffuse within compartments they are anchored to.

**Fig 4.**
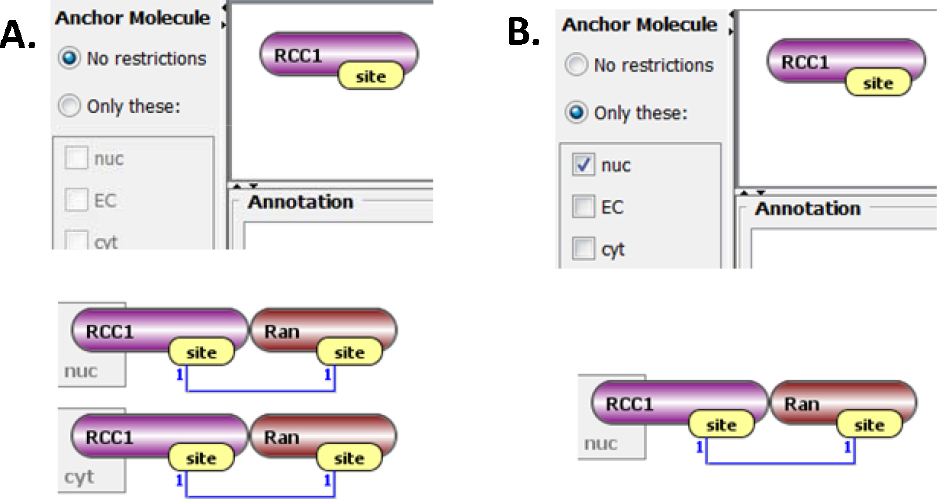
Effect of molecule anchoring. **A**. If “No restrictions” is chosen (top), the rule in Fig 1E will transport RCC1 to the cytosol, producing both species shown at the bottom. **B**. Anchoring will prevent this interaction and the only generated Ran-RCC1 species will be in the nucleus.

As summarized in the introduction, VCell has a hierarchical architecture whereby the system ‘Physiology’ (exemplified by the network in Fig. 3 and all its underlying details) can be associated with several Applications. Applications are used to specify initial conditions, geometries and the physical and mathematical approaches by which the system should be simulated. New Application types have been developed to provide control for network generation and to support a network-free simulations with NFSim. All the simulation methods previously available to manually constructed reaction networks are thereby now available to networks generated automatically from rules. Fig. 5 shows how the Physiology summarized in Fig. 3 can be used to produce compartmental deterministic (ODE), compartmental stochastic, compartmental network-free, spatial deterministic (PDE), spatial stochastic, and spatial hybrid deterministic/stochastic simulations. We believe that VCell is unique in making all these approaches available within one unified software environment; we are also unaware of any other systems biology software offering a spatial hybrid deterministic/stochastic solver(9).

**Fig 5.**
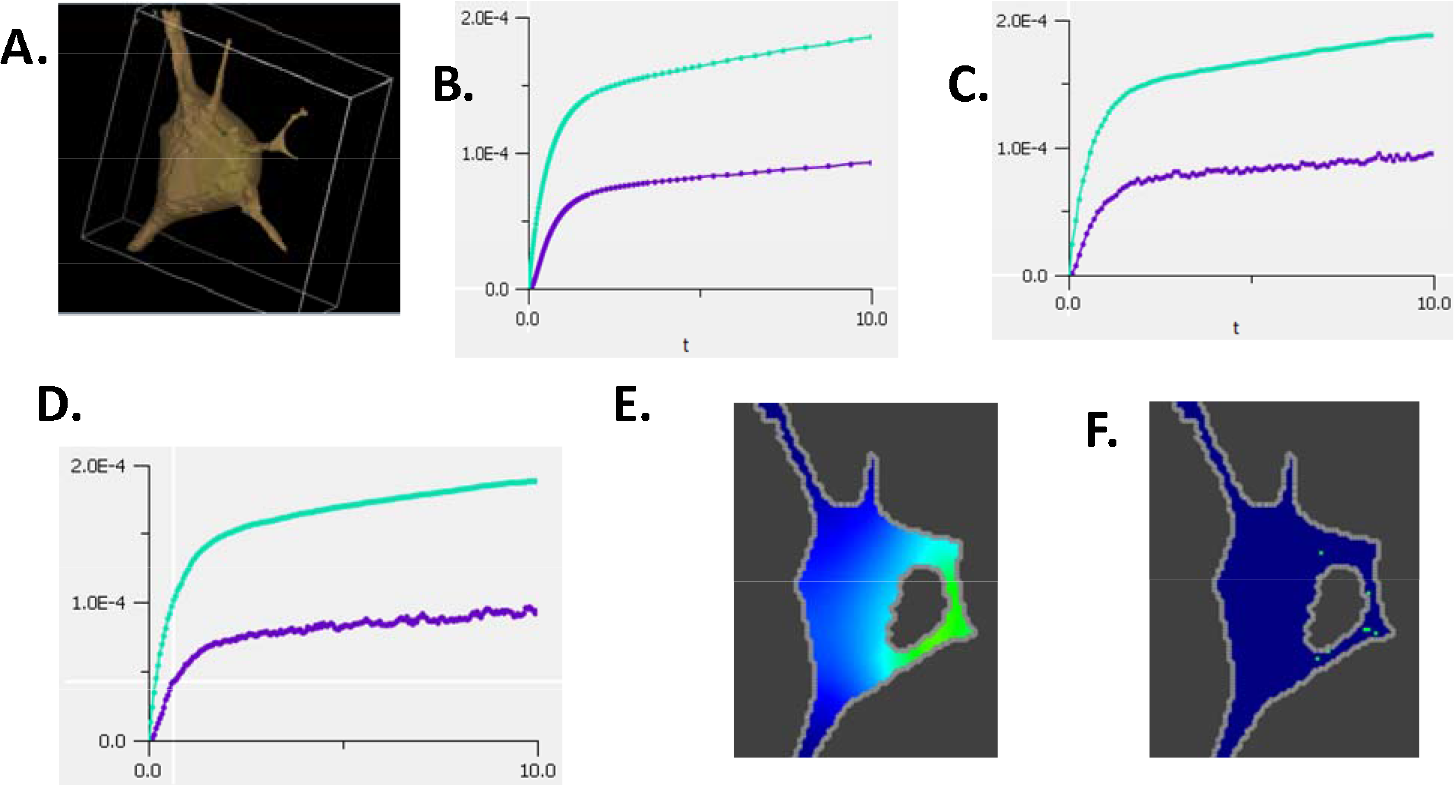
Visualization and simulation capabilities of VCell. **A**. An example of a 3D image-based geometry (neuroblastoma cell) that can be imported into VCell from a confocal microscope. A 3D segmentation within VCell identifies regions corresponding to cytoplasm (tan) and nucleus (green). **B-D**. Simulation results for timecourse of cargo concentration in cytosol of total (upper curve) and phosphorylated (bottom curve) cargo for deterministic (B), stochastic (C), and network-free (D) simulations. **E**. PDE simulation result for one slice displaying the cytosolic cargo concentration gradient at 1 second. Brighter colors correspond to higher concentration. See also Supplementary Movie S1. **F**. Same slice for spatial stochastic simulations for 1,000 molecules of Ran. See also Supplementary Movie S1.

But VCell does not have pretensions of being a solution for every modeling problem. Accordingly, we have devoted significant effort to interoperability – in particular through the SBML standard (15). Specifically for the case of rule-based modeling, a model defined in VCell can be exported to cBNGL (13) for simulation with the stand-alone BioNetGen engine. The generated cBNGL file has a new “anchors block” specifying molecules anchored to compartments. This block is ignored by the BioNetGen compiler. The exported cBNGL has no information about enclosing compartments in the compartment block. Thus, to simulate an exported model with the stand-alone BioNetGen engine, a modeler needs to specify a compartmental tree manually. Also, a model specified in cBNGL can be imported into VCell. Not every file can be seamlessly imported; errors will be displayed when compartment specification is done at a level of individual molecules in seed species and patterns. VCell provides a BNGL import editor where all inconsistencies are displayed and explained, so an experienced BioNetGen user should be able to fix all issues during the import process. Supplemental material S2.pdf provides more details on comparison between cBNGL and VCell representation.

## Discussion

We have described a major enhancement of the VCell software to enable rule-based modeling in multiple compartments. This enhancement gives users with combinatorialy complex biochemical systems the ability to specify all interactions and their dependencies in terms of molecular features such as cellular locations, sites for binding, modification states or conformations. To achieve this, we built a rich GUI that also serves to help visualize the details of these complex systems (Fig. 3). The VCell “classic” manual network generation functionality and GUI are still available and the implementation actually supports mixing of automatically generated rule-based networks with reaction networks generated manually. Such networks can then be modeled with all the VCell compartmental and spatial simulation methods (Fig. 5). For network-free simulations, we have also modified the NFSim engine to support compartments.

There are limitations to rule-based modeling that users should appreciate. One is the restriction of reaction kinetics to mass-action; however, the VCell user may be able to overcome this restriction for deterministic models by judiciously mixing rules and explicit network reactions. A second restriction is that the network-free simulations can only be run for non-spatial models. Additionally, for spatial models based on reaction networks generated by rules, care needs to be exercised not to allow a combinatorial explosion of species and reactions; the resultant large system of PDEs could produce prohibitively expensive computations.

Several rule-based modeling tools that can operate in multiple compartments or perform spatial simulations are available: Simmune (16, 17), KaSim (18), Smoldyn (8), Meredys (19), SRSim (20), SSC (21), SpringSaLaD (22). All of them are exclusively stochastic, whereas VCell offers deterministic spatial and hybrid deterministic/stochastic simulation capabilities. The VCell spatial stochastic solver is based on Smoldyn (23), adapted to permit users to incorporate experimental 3D image-based geometries in simulations; analytical geometries and constructive geometries can also be used. A variety of non-spatial compartmental simulators, both stochastic and deterministic, are also in VCell for quick answers when diffusion is fast on the timescale of reaction kinetics. Additionally, only Simmune and SpringSaLaD have biology-oriented GUIs, as in VCell, while all the other simulators are based on scripting. But the other more specialized simulators have important strengths that might be needed for certain classes of problems. SSC and Smoldyn both have implementations to employ high performance computing or gpus for computationally intensive simulations. Simmune is specialized for complex signaling in immunology. Meredys, SRSim and SpringSaLaD are all designed to account for molecular excluded volume effects and are therefore well suited for simulations where molecular crowding might be important. To facilitate interoperability with such other simulators, VCell supports the SBML standard (15) by enabling export and import of SBML models; it also supports cBNGL export, although some manual editing will be required.

Model sharing is also facilitated within VCell through the VCell database. All models can be stored in the database along with simulations results that were run on the VCell server farm (although users may opt to save models and run simulations on their local machines). Access control is implemented to permit sharing of models with individual collaborators or to make a model openly accessible. Users may annotate model components to connect them to the primary literature sources as well as to ontologies and pathway databases. This is particularly valuable for molecules in rule-based models, where the localization and sites within a molecule can be directly related to both molecular structure and pathway data. Importantly, proper annotation can assure reusability of not just the entire model, but the individual molecules and rules.

## Software availability

Available as VCell (versions 6.1 and later) at the Virtual Cell web site (http://vcell.org/). The application installs and runs on all major platforms and does not require registration for use on the user's computer. Tutorials are available at the Virtual Cell website and Help is provided within the software. Source code is available at https://sourceforge.net/projects/vcell/

## Acknowledgments

The authors gratefully acknowledge useful discussions with James Faeder at University of Pittsburgh and Ann Cowan at UConn Health. The project is supported by the National Institute of General Medical Science through grants P41 GM103313 and R01 GM095485.

## Supporting Material

Supplemental text 1 describes materials and methods. Supplemental text 2 contains a quick software guide. Supplemental movies describe simulation results for spatial deterministic and stochastic applications.

## References

1. Mayer, B. J., M. L. Blinov, and L. M. Loew. 2009. Molecular machines or pleiomorphic ensembles: signaling complexes revisited. J Biol 8:81.81–81.88.

2. Faeder, J. R., M. L. Blinov, B. Goldstein, and W. S. Hlavacek. 2005. Combinatorial complexity and dynamical restriction of network flows in signal transduction. Syst Biol (Stevenage) 2:5–15.

3. Blinov, M. L., J. R. Faeder, B. Goldstein, and W. S. Hlavacek. 2004. BioNetGen: Software for rule-based modeling of signal transduction based on the interactions of molecular domains. Bioinformatics 20.

4. Cowan, A. E., Moraru, II, J. C. Schaff, B. M. Slepchenko, and L. M. Loew. 2012. Spatial modeling of cell signaling networks. Methods Cell Biol 110:195–221.

5. Schaff, J., C. C. Fink, B. Slepchenko, J. H. Carson, and L. M. Loew. 1997. A general computational framework for modeling cellular structure and function. Biophysical journal 73:1135–1146.

6. Schaff, J. C., B. M. Slepchenko, Y. S. Choi, J. Wagner, D. Resasco, and L. M. Loew. 2001. Analysis of nonlinear dynamics on arbitrary geometries with the Virtual Cell. Chaos 11:115–131.

7. Resasco, D. C., F. Gao, F. Morgan, I. L. Novak, J. C. Schaff, and B. M. Slepchenko. 2012. Virtual Cell: computational tools for modeling in cell biology. Wiley Interdiscip Rev Syst Biol Med 4:129–140.

8. Andrews, S. S. 2017. Smoldyn: particle-based simulation with rule-based modeling, improved molecular interaction and a library interface. Bioinformatics 33:710–717.

9. Schaff, J. C., F. Gao, Y. Li, I. L. Novak, and B. M. Slepchenko. 2016. Numerical Approach to Spatial Deterministic-Stochastic Models Arising in Cell Biology. PLoS Comput Biol 12:e1005236.

10. Schaff, J. C., D. Vasilescu, I. I. Moraru, L. M. Loew, and M. L. Blinov. 2016. Rule-based Modeling with Virtual Cell. Bioinformatics 32:2880–2882.

11. Sneddon, M. W., J. R. Faeder, and T. Emonet. 2011. Efficient modeling, simulation and coarse-graining of biological complexity with NFsim. Nat Methods 8:177–183.

12. Faeder, J. R., M. L. Blinov, and W. S. Hlavacek. 2009. Rule-based modeling of biochemical systems with BioNetGen. Methods Mol Biol 500:113–167.

13. Harris, L. A., J. S. Hogg, and J. R. Faeder. 2009. Compartmental rule-based modeling of biochemical systems. In Winter Simulation Conference. 908–919.

14. Smith, A. E., B. M. Slepchenko, J. C. Schaff, L. M. Loew, and I. G. Macara. 2002. Systems analysis of Ran transport. Science 295:488–491.

15. Hucka, M., A. Finney, H. M. Sauro, H. Bolouri, J. C. Doyle, H. Kitano, A. P. Arkin, B. J. Bornstein, D. Bray, A. Cornish-Bowden, A. A. Cuellar, S. Dronov, E. D. Gilles, M. Ginkel, V. Gor, Goryanin, II, W. J. Hedley, T. C. Hodgman, J. H. Hofmeyr, P. J. Hunter, N. S. Juty, J. L. Kasberger, A. Kremling, U. Kummer, N. Le Novere, L. M. Loew, D. Lucio, P. Mendes, E. Minch, E. D. Mjolsness, Y. Nakayama, M. R. Nelson, P. F. Nielsen, T. Sakurada, J. C. Schaff, B. E. Shapiro, T. S. Shimizu, H. D. Spence, J. Stelling, K. Takahashi, M. Tomita, J. Wagner, and J. Wang. 2003. The systems biology markup language (SBML): a medium for representation and exchange of biochemical network models. Bioinformatics 19:524–531.

16. Angermann, B. R., F. Klauschen, A. D. Garcia, T. Prustel, F. Zhang, R. N. Germain, and M. Meier-Schellersheim. 2012. Computational modeling of cellular signaling processes embedded into dynamic spatial contexts. Nat Methods 9:283–289.

17. Zhang, F., B. R. Angermann, and M. Meier-Schellersheim. 2013. The Simmune Modeler visual interface for creating signaling networks based on bi-molecular interactions. Bioinformatics 29:1229–1230.

18. Sorokina, O., A. Sorokin, J. Douglas Armstrong, and V. Danos. 2013. A simulator for spatially extended kappa models. Bioinformatics 29:3105–3106.

19. Tolle, D. P., and N. L. Novère. 2010. Meredys, a multi-compartment reaction-diffusion simulator using multistate realistic molecular complexes. BMC Syst Biol. 4.

20. Gruenert, G., B. Ibrahim, T. Lenser, M. Lohel, T. Hinze, and P. Dittrich. 2010. Rule-based spatial modeling with diffusing, geometrically constrained molecules. BMC Bioinformatics 11.

21. Lis, M., M. N. Artyomov, S. Devadas, and A. K. Chakraborty. 2009. Efficient stochastic simulation of reaction-diffusion processes via direct compilation. Bioinformatics 25:2289–2291.

22. Michalski, P. J., and L. M. Loew. 2016. SpringSaLaD: A Spatial, Particle-Based Biochemical Simulation Platform with Excluded Volume. Biophysical journal 110:523–529.

23. Andrews, S. S., N. J. Addy, R. Brent, and A. P. Arkin. 2010. Detailed simulations of cell biology with Smoldyn 2.1. PLoS computational biology 6:e1000705.

